# Screening the Pathogen Box Compounds for Activity Against *Plasmodium falciparum* Sporozoite Motility

**DOI:** 10.1101/2022.03.30.486498

**Authors:** Sachie Kanatani, Rubayet Elahi, Sukanat Kanchanabhogin, Natasha Vartek, Abhai K. Tripathi, Sean T. Prigge, Photini Sinnis

## Abstract

As the malaria parasite becomes resistant to every drug that we develop, identification and development of novel drug candidates is essential. Many studies have screened compounds designed to target the clinically important blood stages. However, if we are to shrink the malaria map, new drugs that block transmission of the parasite are needed. Sporozoites are the infective stage of the malaria parasite, transmitted to the mammalian host as mosquitoes probe for blood. Sporozoite motility is critical to their ability to exit the inoculation site and establish infection and drug-like compounds targeting motility are effective in blocking infection in the rodent malaria model. In this study, we established a moderate throughput motility assay for sporozoites of the human malaria parasite *Plasmodium falciparum*, enabling us to screen the 400 drug-like compounds from the Pathogen box provided by Medicines for Malaria Venture for their activity. Compounds exhibiting inhibitory effects on *P. falciparum* sporozoite motility were further assessed against transmission-blocking activity and asexual stage growth. Five compounds had a significant inhibitory effect on *P. falciparum* sporozoite motility at 1 μM concentration and four of these compounds also showed significant inhibition on transmission of *P. falciparum* gametocytes to the mosquito and of these four, three had previously been shown to have inhibitory activity on asexual blood stage parasites. Our findings provide new antimalarial drug candidates that have multi-stage activity.

## Introduction

Malaria is caused by parasites of the genus *Plasmodium*, transmitted to humans by *Anopheles* mosquitoes. *Plasmodium falciparum* is responsible for the majority of malaria-induced deaths with over 400,000 deaths and more than 200 million people affected annually (1). As *P. falciparum* is increasingly becoming resistant to artemisinin-based combination therapies (2), new drugs are essential. Drugs used to treat malaria target the asexual blood stages of the parasite, which are responsible for all clinical manifestations of malaria. The majority of these drugs have no impact on the transmission stages of the malaria parasite, and in some cases have been found to increase transmission to the mosquito host (3–5). Recently there has been some focus on developing drugs that target both blood stages and transmission stages, a goal that would enable us to work towards malaria elimination as we treat clinical malaria cases.

Malaria parasites cycle between mosquito and mammalian hosts. Infection in the mammalian host is initiated when mosquitoes inoculate sporozoites as they probe blood. Sporozoites are actively motile, migrating through the dermis to enter the blood circulation (6), which carries them to the liver. Here they develop into exoerythrocytic stages, which produce thousands of hepatic merozoites that initiate the blood stage of infection. Some blood stage parasites differentiate to gametocytes, which are responsible for transmission back to the mosquito. Upon being ingested during blood feeding, gametocytes develop into gametes, fuse, and ultimately form ookinetes, which migrate across the mosquito gut epithelium and develop into oocysts, where sporozoites are produced. These sporozoites migrate to salivary glands and wait to be inoculated into the next mammalian host. Both transmission to the mammalian host and back to the mosquito are severe bottlenecks for the parasite (7). Thus, targeting these stages with drugs could have significant transmission-blocking potential.

Sporozoites move by gliding motility, a substrate-based form of locomotion that does not involve a change in cell shape and is powered by an actin-myosin motor beneath the sporozoite plasma membrane (8). Gliding motility involves the rapid turnover of surface adhesion sites with secretion of adhesins from the apical end followed by their translocation posteriorly via the actin-myosin motor and their shedding via the activity of a surface rhomboid protease (9, 10). Motility is required for sporozoite exit from the dermal inoculation site and entry into hepatocytes, and thus is an excellent target for intervention. Motility studies have largely been performed using the rodent malaria model (11, 12) since to date, it has been difficult to perform live gliding assays with sporozoites of the human malaria parasite *P. falciparum*. In this study, we developed a moderate-throughput *in vitro P. falciparum* sporozoite motility assay and screened the 400 Pathogen Box compounds available from Medicines for Malaria Venture (MMV), which demonstrate activity against a range of different pathogens, predominantly *Mycobacterium* and two groups of eukaryotic protists, the Apicomplexans and Kinetoplastids. Active compounds were further screened against gametocyte transmission to the mosquito and blood stage parasites to identify multi-stage active compounds (Fig. 1A).

**Figure 1.**
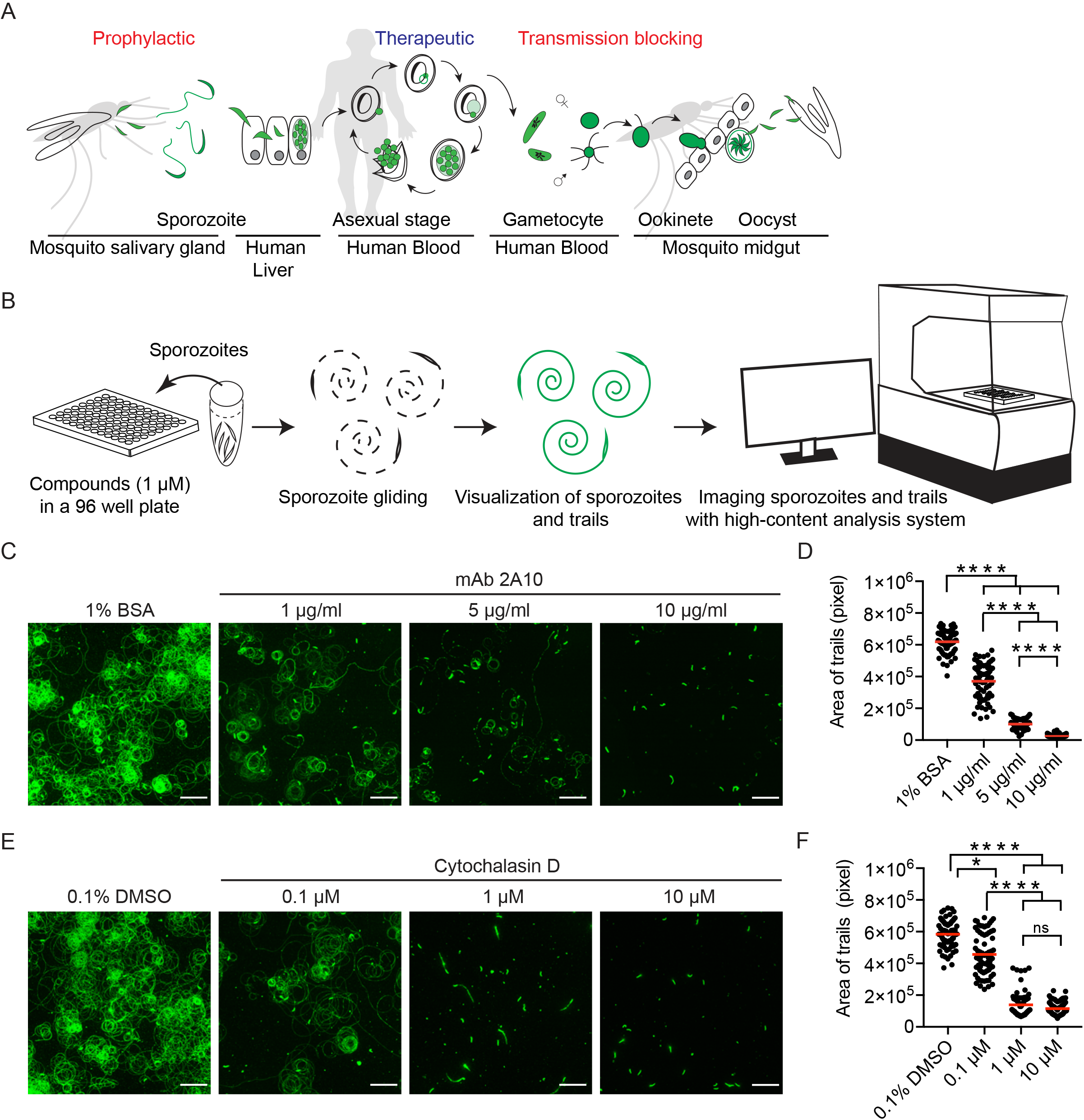
Pathogen box screening strategy and validation. **(A)** Pathogen box screening on different *Plasmodium* life cycle stages. Pathogen box compounds were initially screened for their impact on *P. falciparum* sporozoite motility, which is essential for establishing infection in a human host. Compounds with inhibitory effect on sporozoite motility were further assessed on gametocyte to oocyst development to determine mosquito transmission blocking activity. Finally, compounds with a strong inhibitory effect on both transmission stages were assessed on asexual stage parasite growth. **(B)** Screening strategy of pathogen box compounds on *P. falciparum* sporozoite motility. Freshly isolated *P. falciparum* sporozoites were pre-incubated with pathogen box compounds at 1 μM for 30 minutes, then added to plates and allowed to glide for 1 h in the continued presence of the compound. Trails were visualized by immunofluorescence staining of the circumsporozoite protein (CSP) trails left behind on plates and quantified by high-content imaging of 25 positions in each well. **(C-F)** Validation of the *P. falciparum* motility assay. *P. falciparum* sporozoites were pre-incubated with 0.1 to 10 μg/ml of mAb 2A10 (C&D) or 0.1 to 10 μM Cytochalasin D (E&F) for 30 minutes and added to wells for 1 h in the continued presence of the inhibitor. Sporozoites and trails were then stained for CSP and quantification of the area occupied by fluorescent staining was performed. Panels C&E: Representative images of CSP-stained sporozoites and trails when sporozoites are pre-incubated with the indicated concentrations of mAb 2A10 (C) or Cytochalasin D (E). Scale bars, 50 μm. Panel D&F: Quantification of the area occupied by CSP-stained sporozoites and trails in the indicated treatment groups. Imaging was performed on 25 positions per well with each dot representing the area occupied by fluorescent trails in one image. Data were pooled from 3 independent experiments and all conditions were compared to each other (Kruskal-Wallis test followed by Dunn’s test, **** P < 0.0001, * P < 0.05, ns P > 0.05). Red bars indicate the mean.

## Results

### Moderate throughput *Plasmodium falciparum* sporozoite motility assay

We began by establishing a motility assay for *P. falciparum* sporozoites. We found that if wells of a 96-well plate were coated with mAb 2A10, specific for the repeat region of the sporozoite’s major surface protein, the circumsporozoite protein (CSP), *P. falciparum* sporozoites would adhere to the wells and initiate motility. Unlike rodent malaria sporozoites which will initiate gliding motility on uncoated glass slides, *P. falciparum* sporozoites appear to need the additional adhesive force provided by binding of its surface protein to imobilized antibody in order to glide in two-dimensional spaces. Sporozoites and the CSP trails they leave behind as they move were visualized by staining for CSP, using biotinylated mAb 2A10 followed by avidin conjugated to a fluorophore. Plates were imaged with a high content imaging system and the area occupied by the trails was quantified using cell profiler software (Fig. 1B). This assay was validated by quantification of motility in the presence of known inhibitors, soluble mAb 2A10 (13) and cytochalasin D, an actin polymerization inhibitor (14). Sporozoites were pre-incubated with 1 μM of each compound and then allowed to move in wells of a glass-bottom plate for 1 h. Treatment with either mAb 2A10 or cytochalasin D inhibited sporozoite motility in a dose-dependent manner (Fig. 1C-F), indicating that this assay allows for moderate-throughput measurement of sporozoite motility.

### Screening of pathogen box compounds on *Plasmodium falciparum* sporozoite motility

The pathogen box contains 400 drug-like molecules active against neglected diseases including 125 compounds targeting the blood stages of malaria (https://www.mmv.org/mmv-open/pathogen-box/about-pathogen-box). Screening of the compounds at 1 μM in our sporozoite motility assay demonstrated that five compounds displayed greater than or equal to 50 % inhibition of motility (Sup Fig. 1). We confirmed these results by re-testing the active compounds in 3 biological replicates (Fig. 2A&B). The potency of each compound was then assessed by treating sporozoites with serially diluted compounds from 1 μM to 0.0039 μM. MMV688703 showed the highest potency with significant inhibition at 0.0156 μM and MMV030734 showed second highest potency with significant inhibition at 0.0625 μM (Fig. 2C). The three remaining compounds (MMV688854, MMV687800, and MMV687807) had significant inhibition at 0.25 μM but not at lower concentrations (Fig. 2C). Importantly, the hepatocyte cytotoxicity data provided by MMV showed that 4 of the 5 compounds did not have significant toxicity to HepG2 cells, while MMV687807 showed some toxicity to HepG2 cells (Table 1). We then determined whether these compounds were directly toxic to parasites, using a viability assay based on a live/dead dye that binds to free amines, resulting in dead cells becoming brightly fluorescent (Fig. 3A). Similar to the motility assay, sporozoites were pre-incubated with compounds at 1 μM and the live/dead dye for 30 minutes followed by a 1 h incubation at 37 °C. Sporozoites were then stained for CSP for visualization purposes. Sporozoites treated with any of the 5 motility-inhibiting compounds were > 90 % viable (Fig. 3B), suggesting that these compounds did not have a direct cytotoxic effect on sporozoites.

**Table 1.**
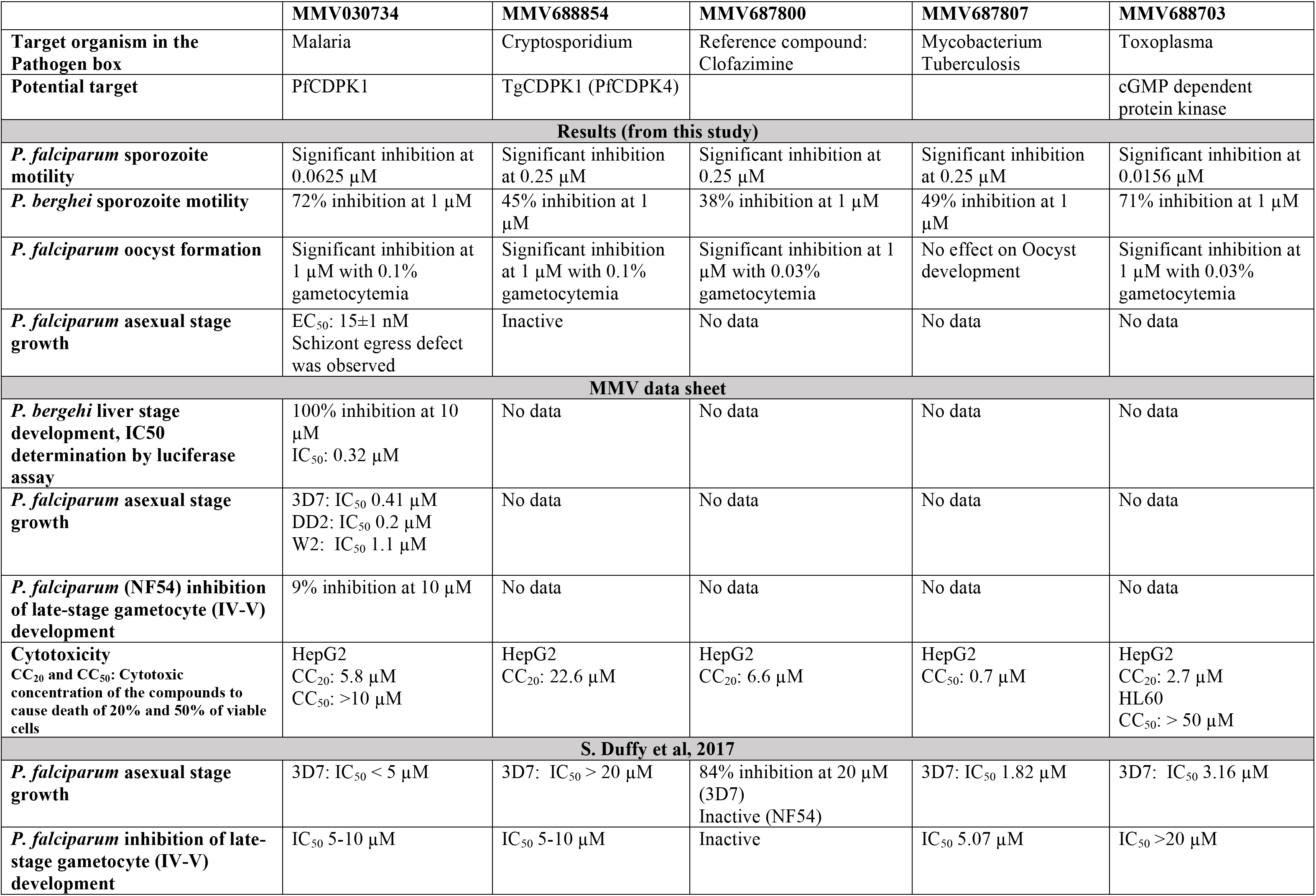
Effect of five compounds in Plasmodium life cycle and potential targets.

|  | **MMV030734** | **MMV688854** | **MMV687800** | **MMV687807** | **MMV688703** |
| --- | --- | --- | --- | --- | --- |
| **Target organism in the Pathogen box** | Malaria | Cryptosporidium | Reference compound: Clofazimine | Mycobacterium Tuberculosis | Toxoplasma |
| **Potential target** | PfCDPK1 | TgCDPK1 (PfCDPK4) |  |  | cGMP dependent protein kinase |
| **Results (from this study)** |  |  |  |  |  |
| ***P. falciparum* sporozoite motility** | Significant inhibition at 0.0625 μM | Significant inhibition at 0.25 μM | Significant inhibition at 0.25 μM | Significant inhibition at 0.25 μM | Significant inhibition at 0.0156 μM |
| ***P. berghei* sporozoite motility** | 72% inhibition at 1 μM | 45% inhibition at 1 μM | 38% inhibition at 1 μM | 49% inhibition at 1 μM | 71% inhibition at 1 μM |
| ***P. falciparum* oocyst formation** | Significant inhibition at 1 μM with 0.1% gametocytemia | Significant inhibition at 1 μM with 0.1% gametocytemia | Significant inhibition at 1 μM with 0.03% gametocytemia | No effect on Oocyst development | Significant inhibition at 1 μM with 0.03% gametocytemia |
| ***P. falciparum* asexual stage growth** | EC<sub>50</sub>: 15±1 nM<br>Schizont egress defect was observed | Inactive | No data | No data | No data |
| **MMV data sheet** |  |  |  |  |  |
| ***P. bergehi* liver stage development, IC50 determination by luciferase assay** | 100% inhibition at 10 μM<br>IC<sub>50</sub>: 0.32 μM | No data | No data | No data | No data |
| ***P. falciparum* asexual stage growth** | 3D7: IC<sub>50</sub> 0.41 μM<br>DD2: IC<sub>50</sub> 0.2 μM<br>W2: IC<sub>50</sub> 1.1 μM | No data | No data | No data | No data |
| ***P. falciparum* (NF54) inhibition of late-stage gametocyte (IV-V) development** | 9% inhibition at 10 μM | No data | No data | No data | No data |
| **Cytotoxicity**<br>CC<sub>20</sub> and CC<sub>50</sub>: Cytotoxic concentration of the compounds to cause death of 20% and 50% of viable cells | HepG2<br>CC<sub>20</sub>: 5.8 μM<br>CC<sub>50</sub>: >10 μM | HepG2<br>CC<sub>20</sub>: 22.6 μM | HepG2<br>CC<sub>20</sub>: 6.6 μM | HepG2<br>CC<sub>50</sub>: 0.7 μM | HepG2<br>CC<sub>20</sub>: 2.7 μM<br>HL60<br>CC<sub>50</sub>: > 50 μM |
| **S. Duffy et al, 2017** |  |  |  |  |  |
| ***P. falciparum* asexual stage growth** | 3D7: IC<sub>50</sub> < 5 μM | 3D7: IC<sub>50</sub> > 20 μM | 84% inhibition at 20 μM (3D7)<br>Inactive (NF54) | 3D7: IC<sub>50</sub> 1.82 μM | 3D7: IC<sub>50</sub> 3.16 μM |
| ***P. falciparum* inhibition of late-stage gametocyte (IV-V) development** | IC<sub>50</sub> 5-10 μM | IC<sub>50</sub> 5-10 μM | Inactive | IC<sub>50</sub> 5.07 μM | IC<sub>50</sub> >20 μM |

**Figure 2.**
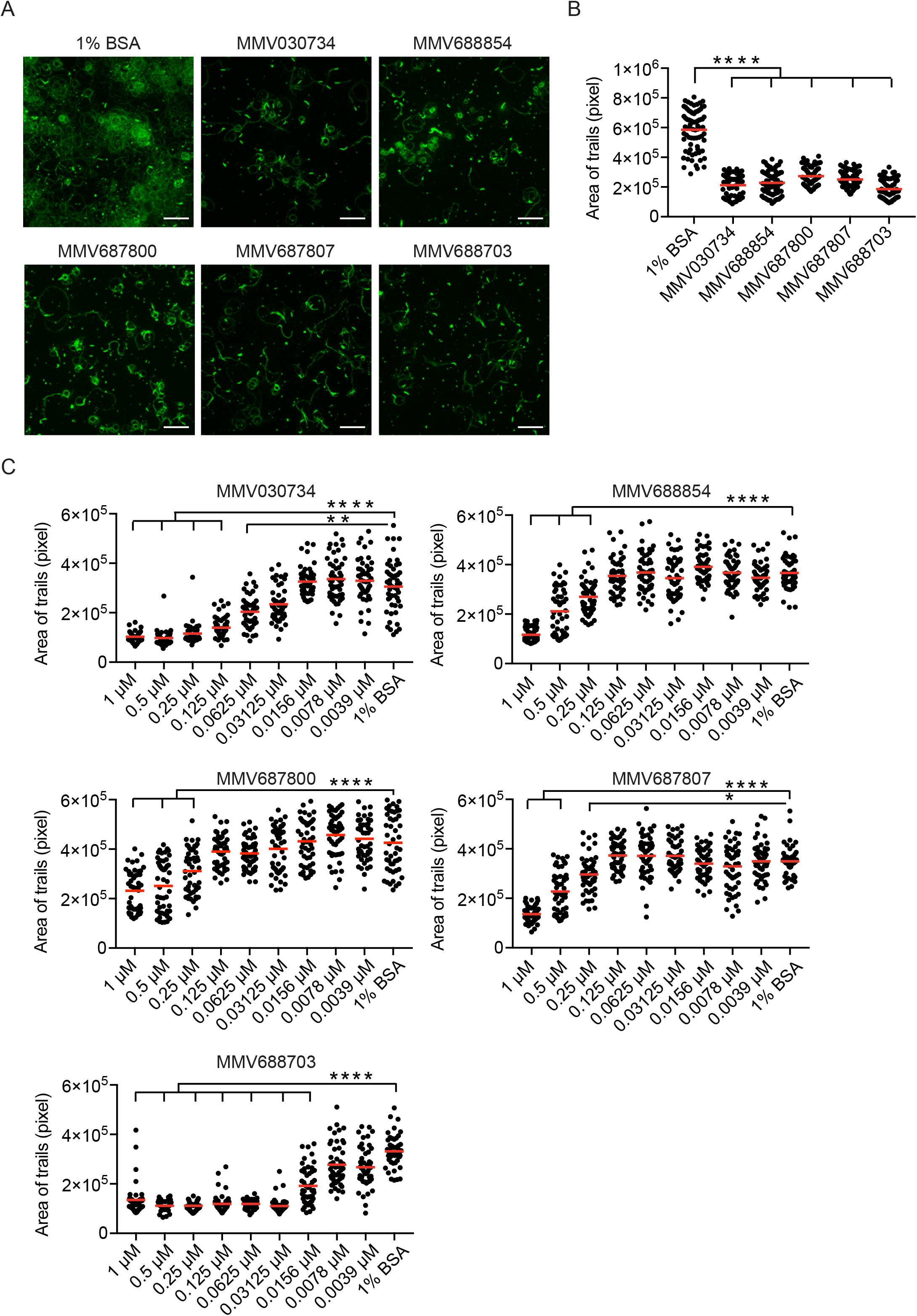
Five pathogen box compounds significantly reduce *P. falciparum* sporozoite motility. *P. falciparum* sporozoites were pre-incubated with the indicated compounds (MMV030734, MMV688854, MMV687800, MMV687807, MMV688703) for 30 minutes and allowed to glide in the continued presence of the compound for 1 h. Following this, sporozoites and trails were stained for CSP and the area occupied by fluorescent sporozoite and trails was quantified by high content imaging. **(A)** Representative images of CSP stained sporozoites and trails in the presence of each of the 5 inhibitory compounds. Scale bars, 50 μm **(B)** Quantification of area occupied by CSP-stained sporozoites and trails in the gliding assays. 25 images per well were analyzed with each dot representing data from one image. Data were pooled from 3 independent experiments and each inhibitor is compared to 1% BSA (Kruskal-Wallis test followed by Dunn’s test, **** P < 0.0001). Red bars indicate the mean. **(C)** Dose-Response of the inhibitory pathogen box compounds. *P. falciparum* sporozoites were incubated with the indicated serially-diluted compounds MMV030734, MMV688854, MMV687800, MMV687807, or MMV688703 and the area occupied by CSP stained trails was quantified. 25 images per well were analyzed with each dot representing data from one image. Data were pooled from 2 independent experiments and serially-diluted compounds were compared to 1% BSA (Kruskal-Wallis test followed by Dunn’s test, **** P < 0.0001, ** P < 0.005, * P < 0.05). Red bars indicate the mean.

**Figure 3.**
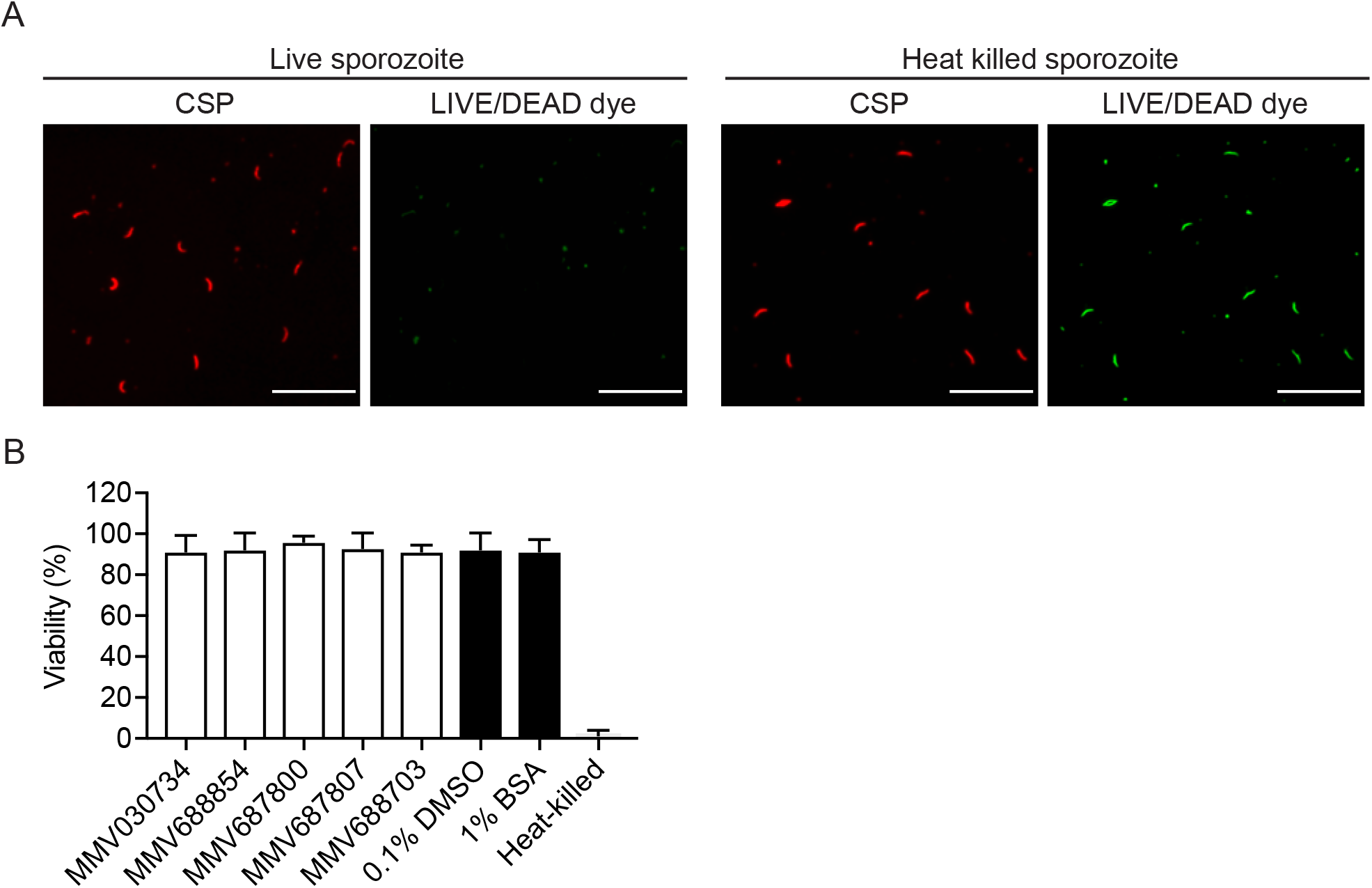
*P. falciparum* sporozoite viability after treatment with each of the five active pathogen box compounds. *P. falciparum* sporozoites were incubated with a fixable live/dead stain and the indicated pathogen box compound for 30 minutes at 20 °C and then moved to 37 °C for 1 h in the continued presence of the stain and compound. Following this, sporozoites were fixed, stained with antibodies specific for CSP, and quantified by immunofluorescence microscopy. **(A)** Representative images of sporozoite stained for CSP (red) and live/dead stain (green). Scale bars, 50 m. **(B)** Quantification of sporozoite viability after treatment with compounds. Total sporozoites (CSP, red) and those that stained with the live/dead stain (green) were counted to determine percent viability (n ≥ 100 sporozoites per condition). Bars representing control samples are shown in black. Shown is the mean +/− standard deviation of two independent experiments.

### Testing the pathogen box inhibitory compounds on *Plasmodium berghei* sporozoite motility

A previous study from the Frischknecht group screened the 400 compounds in the MMV Malaria Box for their impact on sporozoite motility of the rodent malaria parasite *Plasmodium berghei* (11). These compounds are different from those in the MMV Pathogen Box. They found that three of the Malaria Box compounds, MMV665953, MMV665852, and MMV007224, inhibited motility by > 75 %, however, due to toxic effects on hepatocytes, these compounds were not pursued further. To determine if there was cross-species activity of our hits, we tested the active Pathogen Box compounds on *P. berghei* sporozoites. These sporozoites make tighter circles than *P. falciparum* sporozoites complicating quantification of the area occupied by the trails using high content imaging. Thus, we optimized the screening method for *P. berghei* by quantifying stained trails using fluorescence microscopy, measuring total fluorescence intensity using Image J. We verified this assay with mAb 3D11, an inhibitory antibody specific for the *P. berghei* CSP repeats, and cytochalasin D (Fig. 4A&B). As shown, treatment with mAb 3D11 and cytochalasin D inhibited sporozoite motility in a dose-dependent manner. We then assessed the 5 inhibitory Pathogen Box compounds and all of them showed significant inhibitory effect on *P. berghei* sporozoite motility (Fig. 4C&D). MMV030734 and MMV688703 showed > 70 % inhibition while MMV688854, MMV687807, and MMV687800 were less potent. These findings are similar to the inhibitory activity of these compounds on *P. falciparum* sporozoite motility at 1 μM, with MMV030734 and MMV688703 being the most potent and MMV687800 demonstrating the least inhibitory activity (Fig. 2B). Thus, the rodent model can be used for screening compounds targeting motility, though ultimately compounds need to be screened on human malaria parasites.

**Figure 4.**
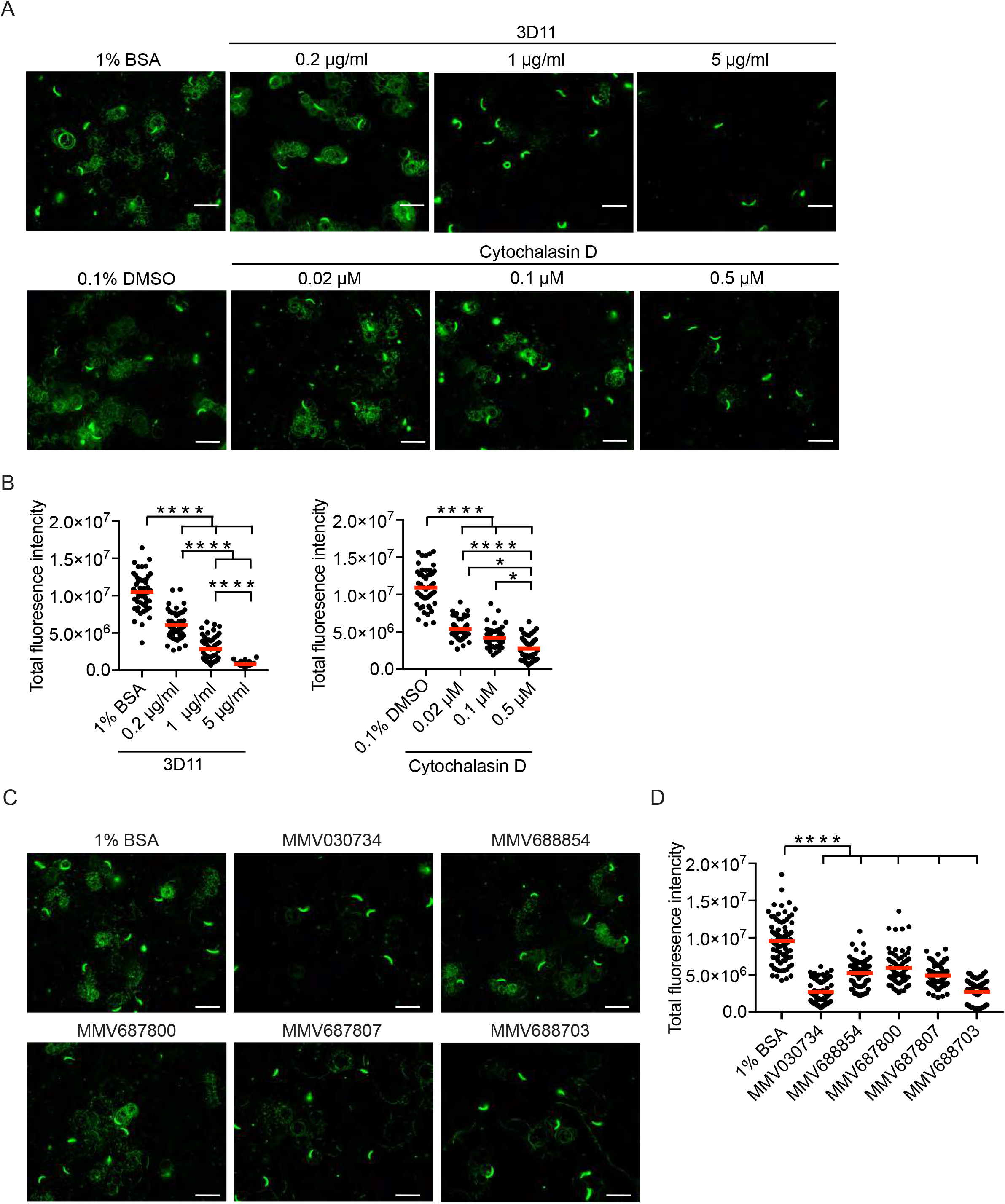
Testing the pathogen box inhibitory compounds on *Plasmodium berghei* sporozoite motility. (A&B) Validation of the *P. berghei* sporozoite motility assay. Sporozoites were pre-incubated with the indicated concentration of mAb 3D11 or Cytochalasin D for 30 minutes and then allowed to glide for 1 h in the continued presence of antibody or Cytochalasin D. Sporozoites and trails were stained for CSP and total fluorescence intensity of sporozoites and trails were quantified by Image J. Representative images of CSP-stained sporozoites and trails are shown in panel A. Scale bars, 20 μm. Panel B shows quantification of the total fluorescence intensity of CSP-stained sporozoites and trails for each treatment group. 25 images were acquired per well with each dot representing fluorescence intensity from one image. Data are pooled from 2 independent experiments and all conditions were compared to each other (Kruskal-Wallis test followed by Dunn’s test, **** P < 0.0001, * P < 0.05). Red bars indicate the mean. **(C&D)** Testing of inhibitory pathogen box compounds on *P. berghei* sporozoites. Sporozoites were pre-incubated with 1 μM of each of the five inhibitory pathogen box compounds (MMV030734, MMV688854, MMV687800, MMV687807, MMV688703) for 30 minutes, then added to plates and allowed to glide for 1 h in the continued presence of the compound. Sporozoites and trails were stained for CSP and fluorescence intensity of sporozoites and trails were quantified by Image J. Representative images of CSP stained sporozoites and trails incubated with the indicated pathogen box compound are shown in panel C. Scale bars, 20 μm. Shown in panel D is quantification of the fluorescence intensity of CSP-stained sporozoites and trails in the presence of the indicated compound. 25 images per well were acquired with each dot representing the fluorescence intensity of one image. Data were pooled from 3 independent experiments and each inhibitor is compared to 1% BSA (Kruskal-Wallis test followed by Dunn’s test, **** P < 0.0001). Red bars indicate the mean.

### Testing the pathogen box inhibitory compounds in transmission blocking assays

We next determined whether any of our active compounds had inhibitory activity on transmission of *Plasmodium* parasites to the mosquito. To do this we added each of the 5 compounds at 1 μM to the *P. falciparum* gametocyte-blood meal fed to mosquitoes. Nine days later, mosquito midguts were dissected and oocyst numbers were counted (Fig. 5A&B). We performed this assay using two different gametocyte concentrations, 0.1% and 0.03% of total erythrocytes. As shown in Fig. 5C, when fed on blood containing low gametocyte counts, there was a significant reduction in oocyst number in mosquitoes fed with four of the compounds, MMV030734, MMV688854, MMV687800, and MMV688703. By contrast, when mosquitoes were fed on blood with higher gametocyte counts, only MMV030734 and MMV68854 had inhibitory activity on transmission. One compound, MMV687807 had no effect on transmission even when blood contained low gametocyte numbers (Fig. 5C and 5D).

**Figure 5.**
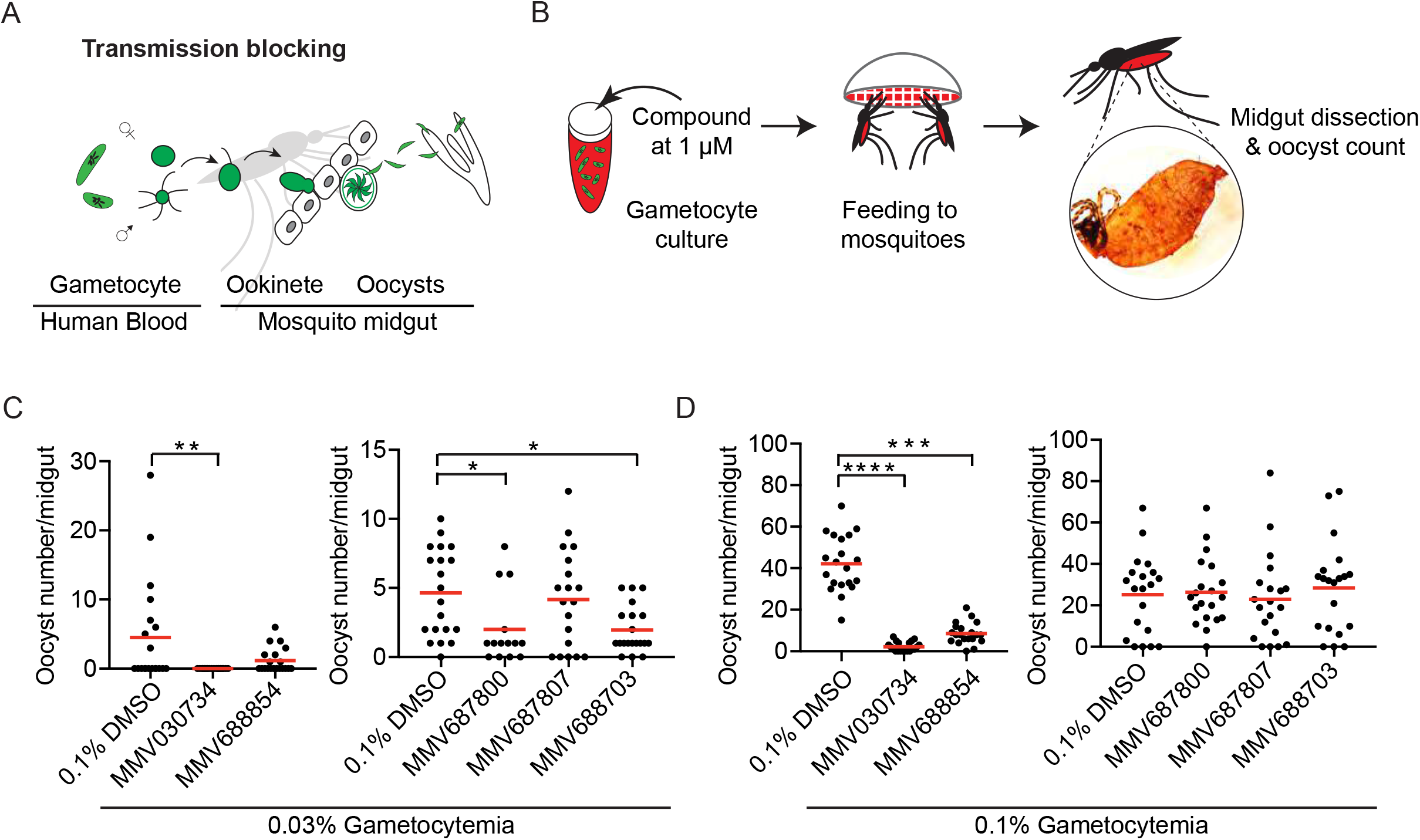
Testing the pathogen box inhibitory compounds for mosquito transmission blocking activity. **(A)** Schematic representation of *Plasmodium* transmission to the mosquito. **(B)** Schematic representation of assay set up. The mature gametocyte culture is mixed with each compound to a final concentration of 1 M and fed to *An. stephensi* mosquitoes. Nine days later, mosquito midguts were dissected and oocysts were counted. **(C&D)** *An. stephensi* mosquitoes were fed a blood meal of either 0.03% (C) or 0.1% (D) gametocytemia in the presence of the indicated compound or DMSO control. On day 8 post-blood meal, individual mosquitoes were dissected and oocysts were counted for 15 to 20 mosquitos per group. Each group is compared to DMSO control (Kruskal-Wallis test followed by Dunn’s test, **** P < 0.0001, *** P < 0.0005, ** P < 0.005, * P < 0.05). Red bars indicate the mean.

### Asexual blood stage parasites treated with MMV030734 exhibit egress defects

Though these compounds had been previously tested against asexual blood stage parasites of *P. falciparum* in a high-throughput assay (15), we wanted to confirm these results for the two compounds (MMV030734 and MMV688854) that had strong inhibitory activity on both transmission stages (Fig. 6A). Synchronized *P. falciparum* ring stage parasites were grown in the presence of 1 μM MMV030734 or MMV688854 or 0.1% DMSO for 60 hours. At the end of the experiment, Giemsa-stained blood smears were made and ring stage parasites were counted. MMV688854 had no impact on parasite growth, however, no ring stage parasites were observed in the culture treated with MMV030734 (Fig. 6B). Interestingly, the Giemsa-stained slides from the MMV030734-treated culture showed that parasite growth was halted at the schizont stage, suggesting an egress defect (Fig. 6C). To confirm this we counted ring, trophozoite, and schizont stage of parasites in infected cells. As shown in Fig. 6D, all parasites were at the schizont stage in the culture treated with MMV030734 while majority of the population in the culture treated with MMV688854 and the DMSO control was ring stage parasites. To further characterize MMV030734, we determined its half-maximal effective concentration (EC_50_) and found that its inhibitory activity on blood stage parasite growth was in the nanomolar range (Fig. 6E).

**Figure 6.**
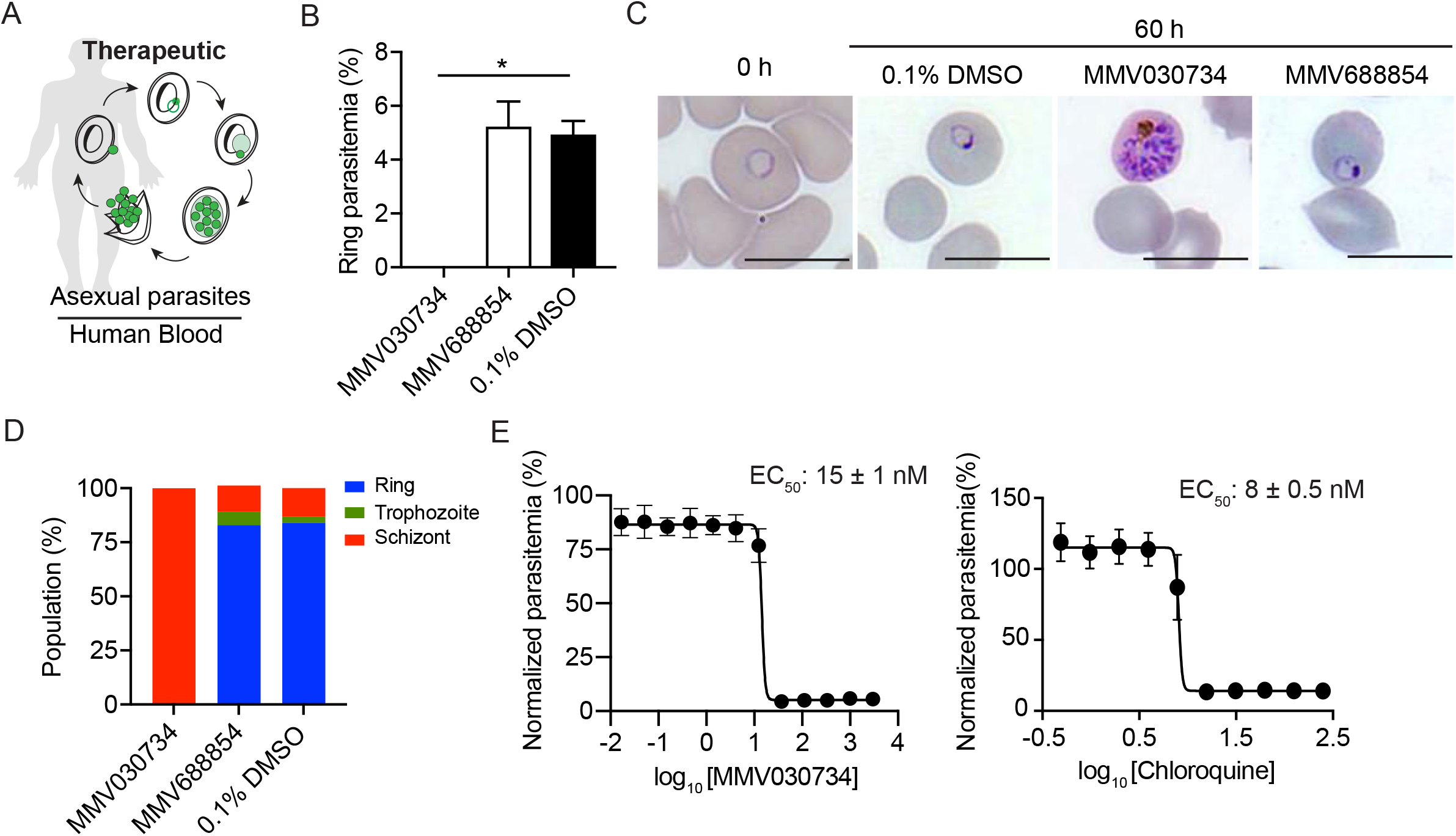
Pathogen box compound MMV030734 reduces asexual stage parasite growth and inhibits egress. **(A)** Schematic representation of *Plasmodium* asexual blood stage cycle. **(B)** Quantification of ring stage parasitemia. Synchronized *P. falciparum* ring stage parasites were incubated with compound MMV030734 or MMV688854 at 1 M or 0.1% DMSO for 60 h at which point Giemsa-stained blood smears were made and ring stage parasites were counted. The mean +/− standard deviation is shown from two technical replicates. ANOVA multiple comparisons, *P<0.01. **(C)**. Representative images of Giemsa-stained asexual stage parasites at 0 h and at 60 h in the indicated treatment group. Scale bar, 10 m. **(D)** Giemsa-stained blood smears taken at 60 h were scored based on the number of ring, trophozoite, and schizont stage parasites. The percentage of each lifecycle stage is shown for one hundred infected erythrocytes from each treatment group. **(E)** Dose-response plot for parasites grown in presence of MMV030734 or Chloroquine for 72 h. Data were pooled from two biological replicates each with four technical replicates. The mean parasitemias ± standard deviation (SD) are shown normalized to the DMSO control. Half maximal effective concentration (EC_50_) is represented as mean ± SD.

**Figure 7.**
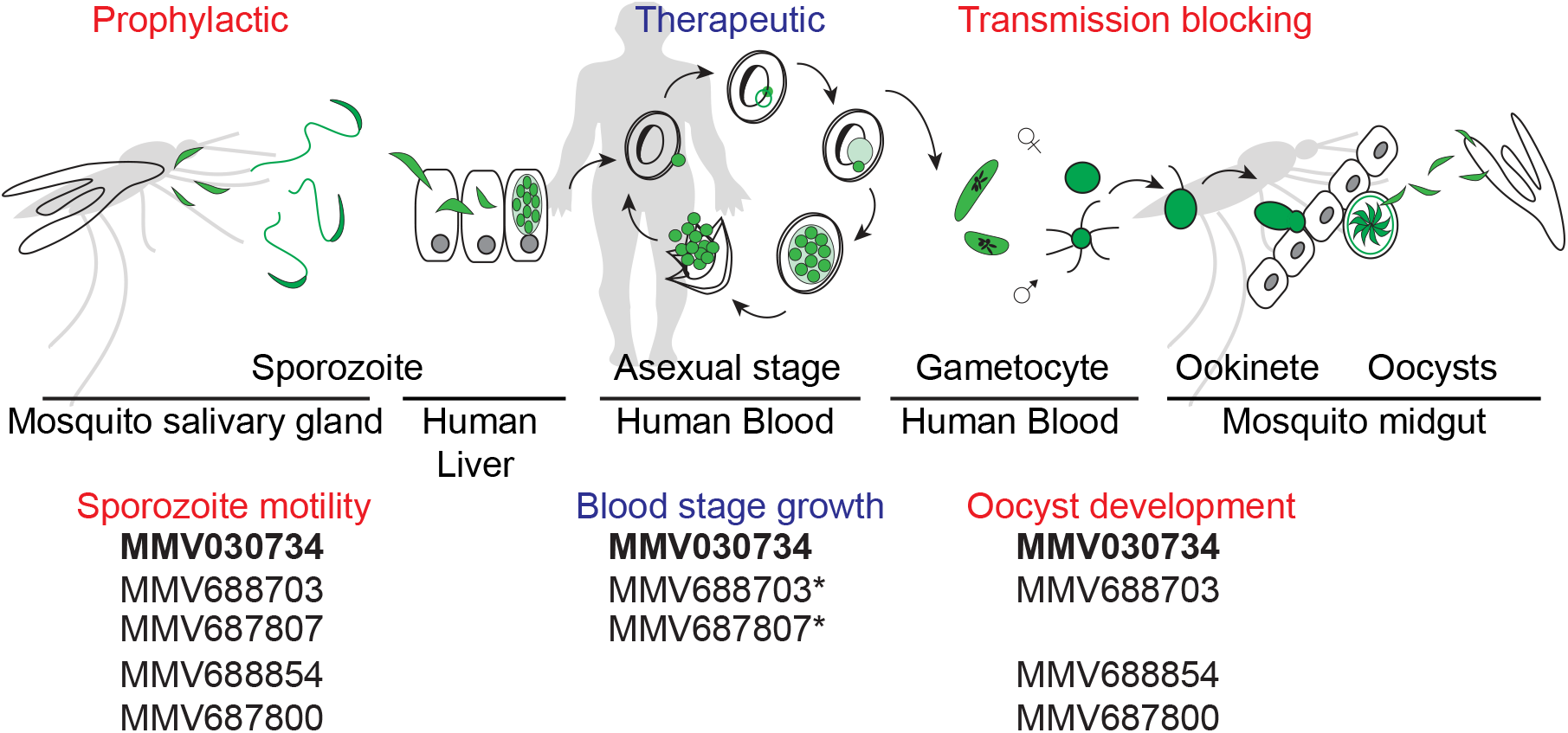
Summary of the five motility-inhibitory compounds across the *Plasmodium* life cycle. Pathogen box compounds were screened for their impact on *P. falciparum* sporozoite motility and 5 compounds (MMV030734, MMV688854, MMV687800, MMV687807, MMV688703) showed significant inhibition in this assay. Of these compounds, MMV030734 and MMV68854 inhibited transmission of *P. falciparum* gametocytes to the mosquito at high gametocytemia and MMV687800 and MMV688703 inhibited transmission only at low gametocytemia. MMV030734, MMV687807 and MMV688703 had a significant inhibitory effect on asexual blood stage growth. Thus, two compounds, MMV030734 and MMV688703 inhibited all three *P. falciparum* life cycle stages. *Blood stage inhibitory effect of MMV688703 and MMV687807 was confirmed by Duffy. S et al (15).

## Discussion

In this study, we have established a quantitative, moderate-throughput assay of *P. falciparum* sporozoite motility and used it to screen 400 drug-like compounds targeting neglected diseases. Five compounds had a significant inhibitory effect on *P. falciparum* sporozoite motility: MMV688854, MMV687800, MMV687807, MMV688703, and MMV030734 (Sup Fig. 1 and Fig. 2). These 5 compounds were further assessed in *P. falciparum* transmission blocking assays and two of them, MMV030734 and MMV688854, had strong inhibitory activity on transmission to the mosquito while two others, MMV687800 and MMV688703, had moderate activity in this assay (Fig. 5). Three of the compounds that had dual transmission-blocking activity, MMV030734, MMV687800, MMV688703, had been previously shown to inhibit growth of *P. falciparum* asexual stage parasites (15), with our study extending the findings with MMV030734 demonstrating an effect on egress from infected erythrocytes (Fig. 6). When the 5 motility-inhibitory compounds were tested in motility assays with the rodent malaria parasite *P. berghei*, all 5 compounds demonstrated inhibitory activity, with MMV030734 and MMV688703 having the greatest inhibitory activity in both species, highlighting the conservation of the gliding motility machinery across the genus, and the usefulness of the rodent model in testing compounds targeting motility.

Interestingly, three of the compounds, MMV688703, MV030734, and MMV688854, are known to target protein kinases (Table 1). MMV688703, a substituted pyrrole, is also known as Compound 1, a cGMP-dependent protein kinase (PKG) inhibitor in *Toxoplasma gondii and Eimeria tenella* (16, 17), MMV030734, a trisubstituted imidazole, binds to *P. falciparum* calcium-dependent protein kinase 1 (PfCDPK1) (18) and inhibits blood stage parasites, and MMV688854, a pyrazolopyrimidine bumped kinase inhibitor derivative, is a known inhibitor of *Toxoplasma gondii* CDPK1 (TgCDPK1) (19, 20). The other compounds that had inhibitory effect on sporozoite motility were MMV687800 and MMV687807, both of which have anti-mycobacterial activity. MMV687800 is Clofazimine, which is used together with dapsone to treat leprosy. Though its precise mechanism of action is unclear, clofazimine interacts with bacterial membrane phospholipids and interferes with K+ uptake and ATP production (21). MMV687807 is a salicylanilide-derivative with significant cytotoxicity (22). There are also 26 reference compounds in the pathogen box and of these, doxycycline, primaquine, amphotericin B, and bedaquiline had significant inhibitory activity against asexual blood stages (15), however they had no impact on sporozoite motility (Sup Fig. 1).

Among the five compounds that inhibited sporozoite motility, MMV688703 or Compound 1, had the greatest potency, with activity in the low nanomolar range. This inhibitor also had activity in blocking parasite transmission to the mosquito. The compound’s target molecule, PKG, is likely the sole mediator of cGMP signaling in *Plasmodium* parasites with phosphoproteomic studies showing hundreds of downstream substrates (23 and reviewed in 22). Thus it is not surprising that PKG regulates many cellular processes in *Plasmodium*, including egress from red blood cells, gamete formation, and motility (25–28). The critical role of PKG in cellular pathways occurring across the life cycle in both mosquito and mammalian hosts, can be explained by the recent demonstration that PKG controls cytosolic Ca^2+^ levels which in turn regulate a variety of stage-specific effector pathways. Its role in motility has been demonstrated in *P. berghei*, where its been shown to inhibit the regulated secretion of micronemes, a calcium-dependent process necessary for gliding motility, and phosphorylation of central components of the actin-myosin motor (28, 29). Our current findings demonstrate that PKG signaling is critical for motility in *P. falciparum*. Additionally, the modest but significant transmission blocking activity of MMV688703 extends previous findings with this compound on gametogenesis and ookinete motility to show that these defects impact mosquito transmission. Taken together these data, covering a wide range of phenotypic assays in both human and rodent malaria parasites, make a compelling case that targeting PKG could have potent multi-stage activity.

Interestingly, the two compounds that had strong transmission-blocking activity in addition to their inhibitory activity on sporozoite motility were MMV688854 and MMV030734, which target kinases of the CDPK family (18–20). CDPKs are serine/threonine protein kinases that are mediators of calcium signaling in *Plasmodium* and other Apicomplexa (30, 31), containing calcium-binding EF hand domains that when bound to Ca^2+^ led to conformational changes, which enable them to rapidly respond to Ca^2+^ fluxes. Apicomplexan parasites have multiple CDPK family members, with *P. falciparum* having 7 CDPKs and the rodent malaria parasites having orthologs to all but one of these. Toxoplasma CDPK1, the target of MMV688854, has been shown to regulate microneme secretion and inhibition of TgCDPK1 results in blockade of parasite motility, host cell invasion, and egress from host cells (32). The *Plasmodium* ortholog of TgCDPK1 is CDPK4. Studies with *P. berghei* demonstrated a role for CDPK4 in sporozoite gliding motility and hepatocyte invasion (29, 33) and in both *P. berghei* and *P. falciparum*, CDPK4 controls microgametocyte activation and exflagellation (34–36). MMV030734 has been shown to target PfCDPK1 which is expressed throughout the *Plasmodium* life cycle, phosphorylates motor complex proteins such as myosin A tail domain-interacting protein (MTIP) and glideosome associated protein 45 (GAP45) (37) consistent with the recent demonstration in *P. berghei* that it plays a critical role in sporozoite motility and invasion (33).

Furthermore, deletion of PfCDPK1 results in slower growth of asexual blood stages and the formation of gametocytes that are not infectious to mosquitoes (38). Recently, the trisubstituted imidazole MMV030084, which is an analogue of MMV030734, has been identified as multi-stage targeting compound, inhibiting *P. berghei* liver stage, *P. falciparum* asexual blood stage development, and male gamete activity (39). Using a chemoproteomics approach, they found that MMV030084 targeted both PfCDPK1 and PKG, however, conditional knockdown and molecular modeling studies pointed to PKG as being the primary target (39).

Our finding that three of the dual-transmission blocking compounds target either PKG or the CDPK family of kinases highlight the central role of calcium signaling as *Plasmodium* parasites move between their mammalian and mosquito hosts and support the idea that PKG, the master regulator of parasite Ca^2+^ levels, and the CDPKs, the effectors of calcium signaling (24, 29, 40), are excellent targets for multi-stage inhibitory drugs. Our new moderate-throughput screening strategy for sporozoite motility facilitates compound testing on *P. falciparum* pre-erythrocytic stages and future work on identifying transmission-blocking compounds and pre-erythrocytic stage vaccine candidates.

## Materials and Methods

### Ethics statement

All animal work was conducted in accordance with the recommendations in the Guide for the Care and Use of Laboratory Animals of the National Institutes of Health. The protocol was approved by the Johns Hopkins University Animal Care and Use Committee (Protocol #M017H325), which is fully accredited by Association for Assessment and Accreditation of Laboratory Animal Care.

### Mosquito infection with *P. falciparum* NF54

Mosquito infection with *P. falciparum* NF54 was performed as previously described (41). Asexual cultures were maintained *in vitro* in O^+^ erythrocytes at 4% hematocrit in RPMI 1640 (Corning) supplemented with 74 μM hypoxanthine (Sigma), 0.21% sodium bicarbonate (Sigma), and 10% v/v heat inactivated human serum. Cultures were maintained at 37°C in a candle jar made from glass desiccators. Gametocyte cultures were initiated at 0.5% parasitemia and at 4% hematocrit. Medium was changed daily for up to 15 to 18 days without the addition of fresh blood to promote gametocytogenesis. Adult *Anopheles stephensi* mosquitoes (3-7 days post-emergence) were allowed to feed through a glass membrane feeder for up to 30 minutes on gametocyte cultures in 40% hematocrit containing fresh O^+^ human serum and O^+^ erythrocytes. Infected mosquitoes were maintained for up to 19 days at 25°C and 80% humidity and were provided with 10% sucrose solution.

### Mosquito infection with *P. berghei* WT-ANKA

Mosquito infection with *P. berghei* WT-ANKA was performed as previously described (42). Swiss Webster mice (Taconic) were infected with *P. berghei* ANKA wild-type parasites and once abundant gametocyte stage parasites were observed, *An. stephensi* mosquitoes (3-7 days post-emergence) were allowed to feed on infected mice. Infected mosquitoes were maintained for up to 25 days at 18°C and 80% humidity and were provided with 10% sucrose solution.

### Pathogen box compounds

Pathogen box compounds were obtained from the Medicines for Malaria Venture (MMV) and consisted of 400 compounds at 10 mM concentration, dissolved in dimethyl sulfoxide (DMSO, Sigma). Compounds were diluted to 1 mM in DMSO and aliquoted into 96 well storage microplates (Sigma, CLS3363) and stored at −80°C.

### *P. falciparum* moderate-throughput sporozoite motility assay

15,000 freshly dissected *P. falciparum* salivary gland sporozoites in 60 μl of 2% bovine serum albumin (BSA) in Hank’s Balanced Salt Solution (HBSS) at pH 7.4 were mixed with 60 μl of each pathogen box compound at 2 μM which and added to a well of a U-bottom 96 well plate (Falcon, 353077). The final concentration of each pathogen box compound was at 1 μM in 1% BSA in HBSS. The plate was incubated for 30 minutes at 20°C and sporozoites and compound mixture were transferred to a 96 well glass bottom plate (Griner, 655892) coated with 5 μg/ml of mAb 2A10 in PBS. The plate was centrifuged for 3 minutes at 300 × g and incubated for 1 h at 37°C. Wells were fixed in 4% paraformaldehyde in PBS, blocked with 1% BSA in PBS (pH 7.4) and stained with biotinylated mAb 2A10 in 1% BSA in PBS (pH 7.4) for 1 h at room temperature, followed by detection with Alexa Fluor 488 streptavidin (Invitrogen, S11223) diluted at 1:500 in PBS for 1 h at room temperature. Samples were preserved in a glycerol / PBS solution (ratio, 9:1) at 4°C and the plate was imaged within one week. Imaging was performed on 25 positions per well (5 x 5, 500 μm apart) by using ImageXpress Micro XLS Widefield high-content analysis system (Molecular Devices) with 40X Plan fluor objective.

### Quantification of area occupied by trails using Cell Profiler software

Image analysis was automated with the open source Cell Profiler software (version 3.0.0) (43). All images were run through a pipeline designed to threshold the images and quantify area occupied by trails. Image intensity was rescaled from 0 - 0.007 to 0 – 1 and the rescaled image was used to set the threshold which was set automatically by using the minimum cross entropy set up in the Cell Profiler pipeline. Following this, object pixel diameter size between 5 to 1,000 was counted and exported to an excel file.

### Testing pathogen box compounds on *P. berghei* sporozoite motility

Coverslips were placed in 24-well plates and coated with 5 μg/ml of mAb 3D11 in PBS for 1 h at room temperature and then washed. A total of 50,000 sporozoites in HBSS with 2% BSA (pH 7.4) were mixed with pathogen box compounds, cytochalasin D or mAb 3D11 in a low retention 1.5 ml tube (Axygen, MCT-175-L-C) and pre-incubated for 30 minutes at 20°C. Each sporozoite and compound mixture was then added to an mAb 3D11-coated well, centrifuged onto the coverslip for 3 minutes at 300 × g and incubated for 1 h at 37°C. Wells were fixed in 4% paraformaldehyde in PBS, blocked in 1% BSA in PBS (pH 7.4) and stained with biotinylated mAb 3D11 diluted at 1:500 in 1% BSA in PBS (pH 7.4) for 1 h at room temperature, followed by detection with Alexa Fluor 488 streptavidin (Invitrogen) diluted at 1:500 in PBS for 1 h at room temperature. Samples were mounted in gold antifade mountant (Invitrogen, P36935) and imaged with fluorescence microscopy (Nikon E600) using 40X objective. Twenty-five positions per slide were acquired using identical exposure settings and acquired images were batch processed using Fiji (https://fiji.sc/) to measure total fluorescence intensity.

### Sporozoite viability assessment

20,000 freshly dissected sporozoites were pre-incubated with pathogen box compounds at 1 μM concentration and 1:1,000 diluted live/dead fixable green stain (Invitrogen, L23101) in HBSS with 1% BSA (pH 7.4) at 20°C for 30 minutes and then transferred to 48 well plate containing a coverslip, centrifuged at 300 × g for 3 minutes and further incubated at 37°C for 1 h, to replicate the treatment of sporozoites in the gliding assay. After incubation, sporozoites were fixed with 4% PFA, blocked with 1% BSA in PBS (pH 7.4) and stained with 1 μg/ml of mAb 2A10 in 1% BSA in PBS (pH 7.4) followed by detection with Alexa fluor 594 goat anti-mouse secondary antibody (Invitrogen, A11032). Samples were mounted in gold antifade mountant (Invitrogen) and observed by fluorescence microscopy. For each condition, 100 sporozoites were identified by CSP staining (red) and dead sporozoites were counted by live/dead stain (green) to quantify viability.

### Testing pathogen box compounds using the standard membrane feeding assay (SMFA)

The pathogen box compounds (MMV030734, MMV688854, MMV687800, MMV687807, MMV688703) or DMSO in HBSS were mixed with gametocyte cultures such that compound final concentration was at 1 μM and DMSO was at 0.1% and fed to adult female *An. stephensi* mosquitoes (3-7 days post-emergence). Cultures with final gametocytemias of 0.3% and 0.01% were used for feeding. *An. stephensi* mosquitoes were allowed to feed for up to 30 minutes. Infected mosquitoes were maintained at 25°C and 80% humidity and were provided with 10% sucrose solution. Oocyst development was quantified on day 9 post blood feeding by staining mosquito midguts with 0.1% mercurochrome (Sigma, M7011) in PBS and counting by bright-field microscopy with a 10X objective.

### Testing pathogen box compounds on *Plasmodium falciparum* asexual stages

The *P. falciparum* transgenic NF54^attB^ line (44) which has similar EC_50_ of chloroquine to NF54 wild type (45, 46) were cultured in O^+^ erythrocytes at 2% hematocrit and maintained in 25 cm^2^ gassed flasks (94% N_2_, 3% O_2_, and 3% CO_2_) at 37°C. The cultures were kept in RPMI 1640 medium with L-glutamine (US Biological, R8999) supplemented with 20 mM HEPES, 0.2% sodium bicarbonate, 12.5 μg/mL hypoxanthine, 5 g/L Albumax II (Life Technologies, S7563), and 25 μg/mL gentamicin. Cultures were synchronized using 5% (wt/vol) sorbitol as previously outlined (47) and synchronized ring parasites were seeded at 1% parasitemia and 1% hematocrit and treated with 1 μM of the indicated pathogen box compounds or 0.1% DMSO or 1 μM chloroquine for 60 hours. After incubation at 37°C for 60 h, parasite growth was quantified by Giemsa-stained blood smear. To determine the half-maximal effective concentration (EC_50_) of MMV030734, sorbitol-synchronized ring parasites were seeded at 3% parasitemia and 2% hematocrit in a 96-well flat bottom plate and grown in the presence of inhibitor at 37°C for 72 h. A concentration series for MMV030734 (17 pM to 3 μM) and chloroquine (487 pM to 250 nM) was tested. After 72 h, parasite growth was quantified using SYBR Green I (Invitrogen) to stain DNA and an Attune NxT Flow Cytometer as described previously (48, 49). Parasitemia from the MMV030734 and chloroquine samples were normalized to the 0.1% DMSO control. Data from two independent biological replicates, each with four technical replicates, were fit to a four-parameter sigmoidal dose-response curve using Prism V8.4 (version 8.4, GraphPad).

### Statistical analysis

All statistical analyses were performed with Graphpad Prism (Version 7 or 8.4).

## Acknowledgements

We would like to thank the team of the parasitology and insectary core facilities at the Johns Hopkins Malaria Research Institute, the Johns Hopkins School of Medicine Microscopy Facility (MicFac) and Hoku West-Foyle for invaluable assistance. This work was supported by the National Institutes of Health (R01 AI132359 to PS, R01 AI065853 to STP), Johns Hopkins Malaria Research Institute postdoctoral fellowships (SK and RE), Bloomberg Family Philanthropies, and NIH grants that funded the MicFac high content imager (R01GM28007-S1 and R01GM66817-S1).

---

**Supplemental Figure 1.**
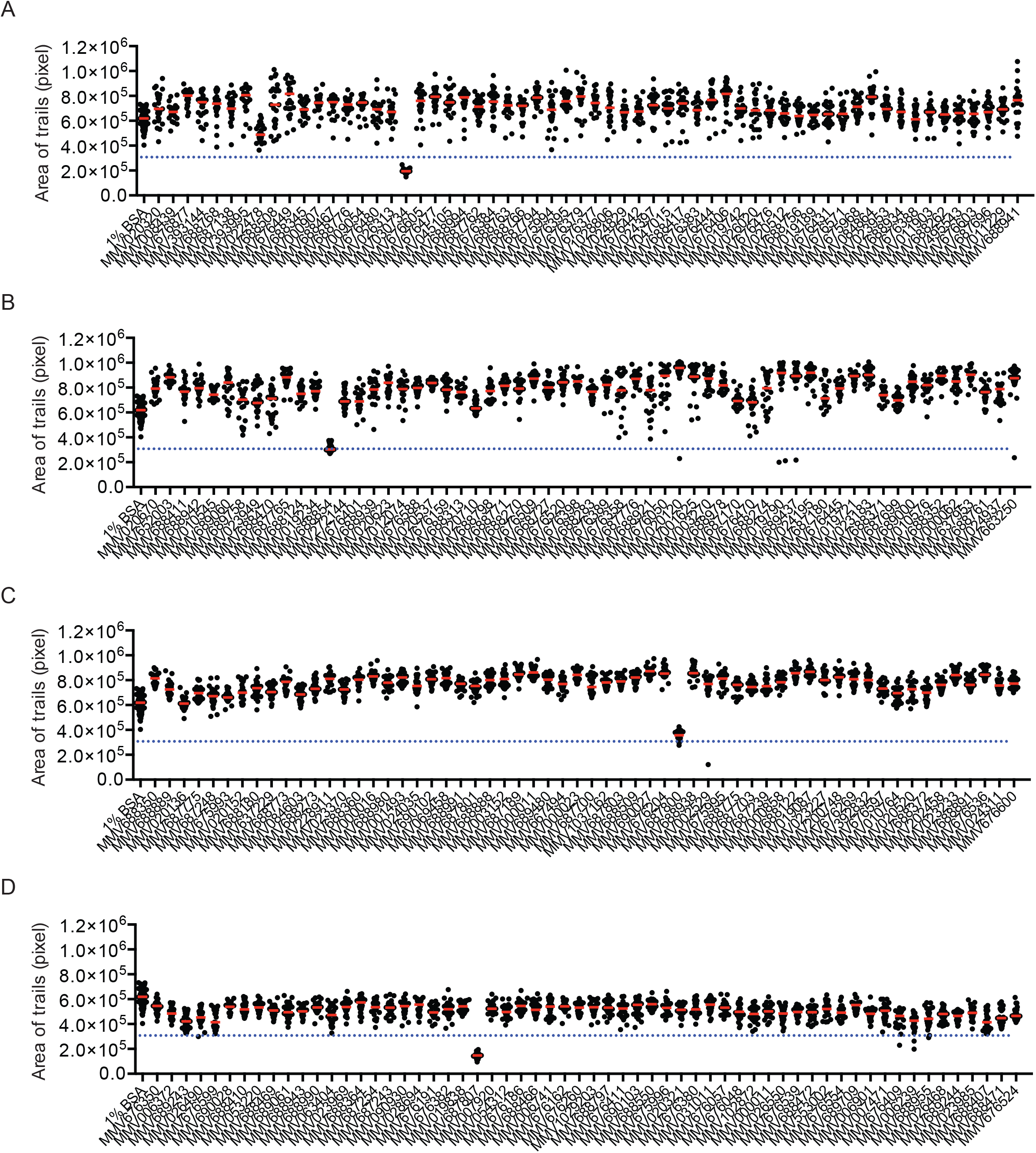
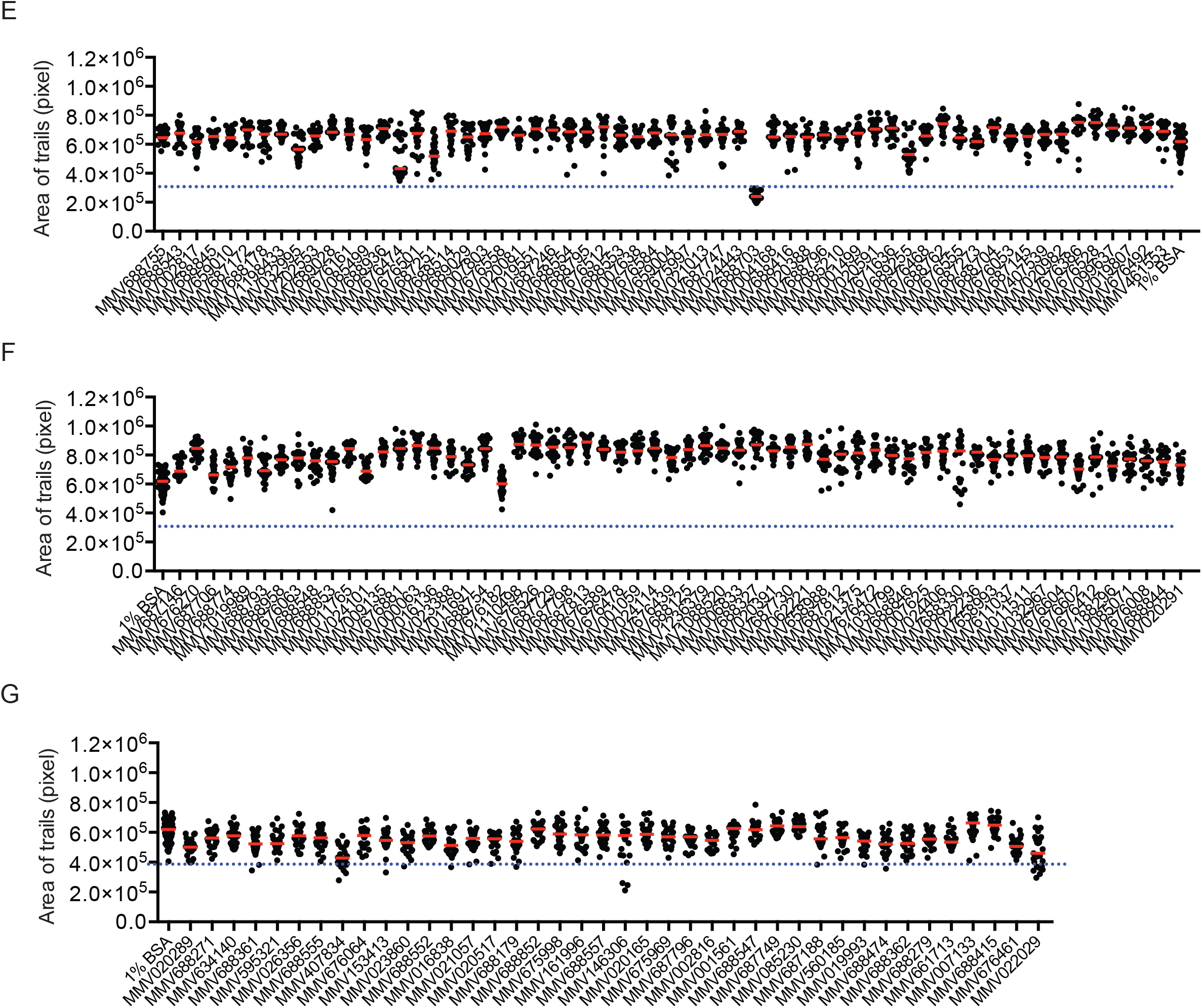
Screening of pathogen box compounds on *P. falciparum* sporozoite motility. Freshly isolated *P. falciparum* sporozoites were incubated with each of the 400 pathogen box compounds at 1 M and allowed to glide for 1 h. CSP trails were stained and quantification of the area occupied by the trails was performed. Red bars indicate the mean area occupied by trails. Each panel shows a different subset of the 400 compounds. Inhibition of motility was observed in sporozoites treated with MMV030734 (panel A), MMV688854 (panel B), MMV687800 (panel C), MMV687807 (panel D), MMV688703 (panel E). The dotted line in each graph indicates the threshold for identifying a compound hit. The threshold is half of the mean area occupied by the trails of parasites in 1% BSA.

